# Sequence dependencies and biophysical features both govern cleavage of diverse cut-sites by HIV protease

**DOI:** 10.1101/2022.04.18.488666

**Authors:** Neha Samant, Gily Nachum, Tenzin Tsepal, Daniel N.A. Bolon

## Abstract

The infectivity of HIV-1 requires its protease cleave multiple cut-sites with low sequence similarity. The diversity of cleavage sites has made it challenging to investigate the underlying sequence properties that determine binding and turnover of substrates by PR. We engineered a mutational scanning approach utilizing yeast display, flow cytometry, and deep sequencing to systematically measure the impacts of all individual amino acid changes at 12 positions in three different cut-sites (MA/CA, NC/p1, and p1/p6). The resulting fitness landscapes revealed common physical features that underlie cutting of all three cut-sites at the amino acid positions closest to the scissile bond. In contrast, positions more than two amino acids away from the scissile bond exhibited a strong dependence on the sequence background of the rest of the cut-site. We observed multiple amino acid changes in cut-sites that led to faster cleavage rates, including a preference for negative charge five and six amino acids away from the scissile bond at locations where the surface of protease is positively charged. Analysis of individual cut sites using full-length matrix-capsid proteins indicate that long-distance sequence context can contribute to cutting efficiency such that analyses of peptides or shorter engineered constructs including those in this work should be considered carefully. This work provides a framework for understanding how diverse substrates interact with HIV-1 protease and can be extended to investigate other viral proteases with similar properties.

## Introduction

The interdependence of mutations, also known as epistasis, can provide valuable insights into biochemical mechanism and are also critical to understanding evolution. In terms of mechanism, mutational dependencies have revealed concerted motions in proteins that govern allostery and protein evolution [1, 2], provided physical maps to improve the prediction of protein structure [3, 4], and identified mutations that increase thermodynamic stability [5]. In evolution, epistasis can have dramatic impacts on the rate and pattern of substitutions [6, 7].

While the value of understanding epistasis is clear, the prevalence, magnitude, and evolutionary impacts of mutational dependencies are controversial [8]. For example, some studies [9] support the idea that the preferences of amino acids at a position in a protein change during evolution whereas, others [10] indicate that key aspects of these preferences remain largely conserved during evolution. Part of the controversy regarding the impacts of epistasis stems from a dependence on the resolution of fitness measurements. According to well established population genetic theory, the magnitude of fitness effects that contribute to selection in natural evolution is approximately the inverse of the effective population size [11]. For example, most microbes exhibit effective population sizes of greater than a million meaning that fitness effects of one in a million would contribute to natural selection. The patterns of epistasis discerned from analyses of sequence patterns in natural evolution will be determined by minute fitness effects. In contrast, epistasis from experimental measures of fitness will be determined by their resolution that is typically on the order of one in a hundred to one in a thousand. Investigation of experimentally assessed epistasis across studies ranging from viruses to animals indicated the greatest level of epistasis in viruses [12]. This study suggests that compact genomes (e.g., as in HIV-1) tend to experience high levels of epistasis. While some of the features contributing to patterns of epistasis in different organisms have been investigated, the structural and physical underpinnings of epistasis remain poorly understood for most proteins and organisms.

Here, we investigate the structural and physical underpinnings of epistasis in cleavage sites of HIV-1 protease (PR). PR cut-sites provide a rich system for investigating epistasis because they exhibit dramatic sequence variation (Figure 1A). PR recognizes and cleaves its substrate Gag and Gag-Pol polyproteins at twelve different sites that are required to generate mature viral proteins and infectious viruses. The twelve cleavage sites are highly variable in amino acid sequence (Figure 1A). Despite the varied sequence of cut-sites, structural studies have shown that they occupy a conserved shape in the PR active site [13-15].

**Figure 1.**
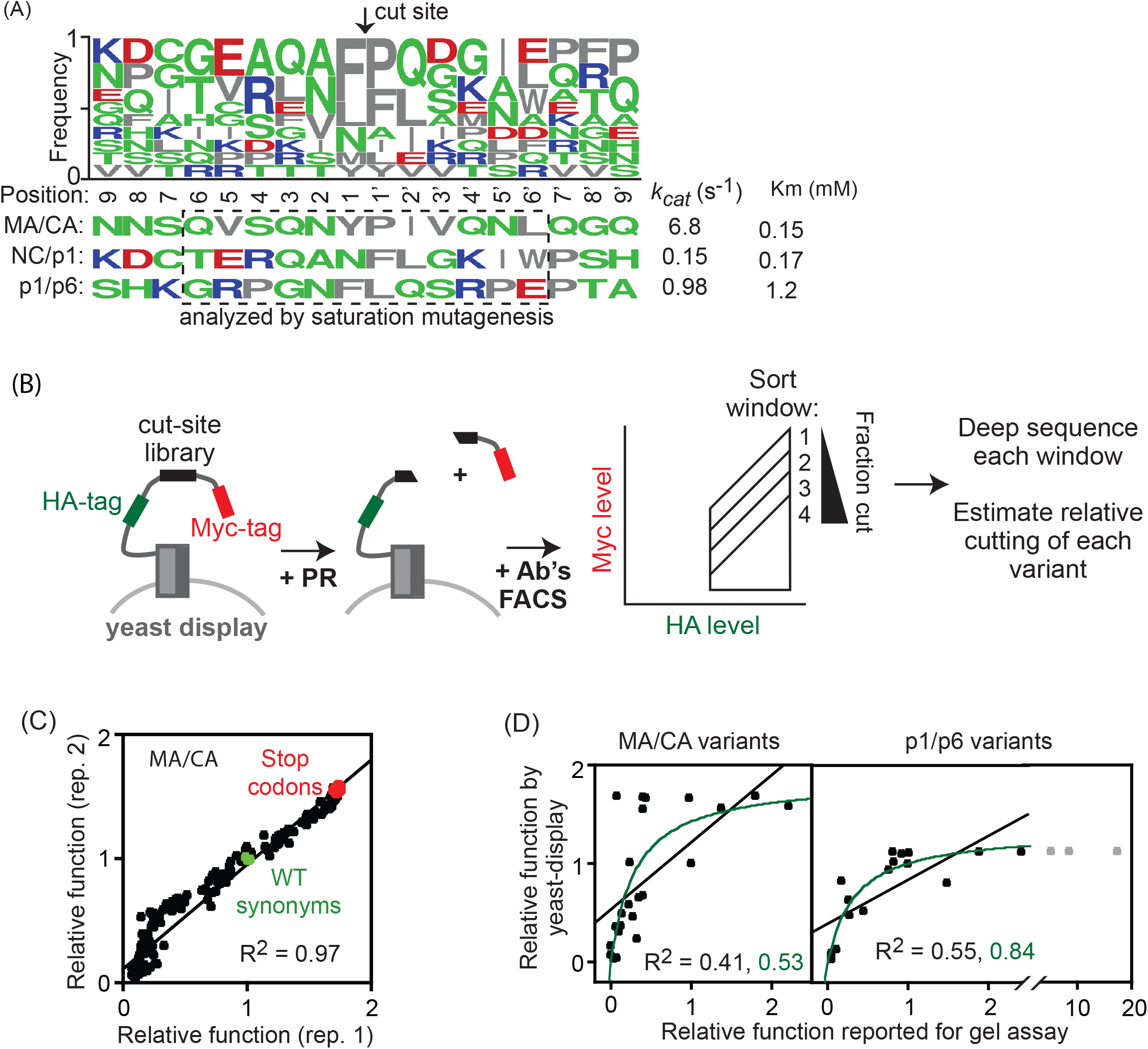
Yeast display approach to examine physical underpinnings of cut-sites for HIV-1 protease. (A) Logo plot showing the frequency of each amino acid in an alignment of all PR cut-sites in the proteome of the NL4-3 HIV-1 strain. The WT cut-sites analyzed in this work (MA/CA, NC/p1 and p1/p6) are shown below. (B) Outline of yeast-display approach to measure the impacts of all point mutations on the function of cut-sites. (C) Correlation between full biological replicates of yeast display measures of functional effects in the MA/CA cut-site. (D) Correlation between yeast display estimates of cutting function and previously reported measures using a gel-based assay for a panel of variants in the MA/CA and p1/p6 sites [40]. Correlations were estimated for both linear fits (shown in black), and non-linear fits (shown in green). The grey points were excluded from fits to avoid potential lever arm effects on linear fits to data points with extreme values.

PR is a 99 amino acid protein that functions as a homodimer with one active site per dimer [14]. A catalytic aspartate at position 25 activates a water molecule for peptide-bond hydrolysis of substrates. The active site surrounding the catalytic aspartate is a groove that directly contacts the four amino acids immediately before and after the cleavage site [14, 16]. The amino acids adjacent to the cut site are referred to as position 4-3-2-1/1’-2’-3’-4 (cut site indicated by /). While positions 4 to 4’ directly contact the active site [14], recent studies indicate that substrate positions beyond P4 and P4’ can contribute to substrate binding [16, 17].

While all cut-sites must be cleaved by PR in order to generate infectious virions, the turnover rate (*k*_*cat*_) of sites vary over a wide range: from 2×10^−5^ sec^-1^ for the NC/TFP site to 7 sec^- 1^ for the MA/CA site [18, 19]. The wide range of substrate cleavage rates led to the suggestion that the order of cleavage may be important for viral maturation [20, 21]. Recent studies [18, 22] indicate that the cutting rate of many sites can be altered dramatically without compromising fitness. However, the relationship between viral fitness and cutting efficiency has not been directly assessed for most cut sites.

Protease inhibitors (PIs) potently inhibit replication of the wildtype HIV-1, and the virus responds through the accumulation of mutations in both PR and some cut-sites [23, 24]. High level resistance to PI requires the accumulation of multiple mutations in PR, typically involving 10-30 amino acid changes in this 99 residue protein [25, 26]. To our knowledge, all PR variants that cause resistance to PI’s exhibit dramatically reduced enzyme activity [27-30]. Many studies have found that cleavage sites in Gag co-evolve with drug resistant mutations in PR, indicating that the mutations in the cleavage sites may compensate for reduced enzyme activity of PR [31-39]. Cut-site compensatory evolution has most commonly been associated with the NC/p1 and p1/p6 sites.

While there is extensive evidence that cut-sites are critical for viral replication and subject to selection pressure by PI’s, the wide variation in cut-site sequences has made it challenging to elucidate the features that underly cut-site recognition and processing by PR. Prior studies have found evidence supporting both context-dependent effects of mutants [17, 40] and context-independent features [41], but it is not clear how differences in experimental approaches may contribute to these conclusions. To address this challenge, we analyzed how all possible single amino acid changes in three different cut sites (MA/CA, NC/p1, and p1/p6) impact cleavage rate by PR in the same assay. The resulting cut-site fitness landscapes reveal both general biophysical features that underlie cutting efficiency in all three cut sites as well as features that depend on the rest of the sequence background of the cleavage site.

## Results and Discussion

### Experimental approach to investigate features underlying cleavage efficiency

The sequence of the different sites that HIV-1 PR must cut in order to generate an infectious virus vary dramatically, which has made it challenging to understand the sequence rules that govern cleavage. The cut-sites encoded by a single viral genome (NL4-3) are shown in Figure 1A. All residues adjacent to the scissile bond vary, with no position showing even 50% sequence identity. The two closest residues on either side of the scissile bond exhibit the least amino acid variation, but even these sites include amino acids that vary dramatically in physical properties including size, hydrophobicity, and hydrogen bond potential. Positions three or more sites from the scissile bond exhibit similar highly diverse amino acids.

To study both general and sequence-dependent features, we chose to analyze the fitness landscape at 12 positions in three different cut-sites (MA/CA, NC/p1, and p1/p6). The sequence of these cut-sites have two identical amino acids within the 12 amino acid region of study (N at position 2 in MA/CA and p1/p6, and Q at position 3 in MA/CA and NC/p1). This level of identity (2 of 36 comparisons = 5.5%) closely matches expectations for comparisons of random amino acid sequences (1/20 = 5%). We were motivated to investigate features of NC/p1 and p1/p6 that may underlie their observed co-evolution with drug-resistant mutations in PR. In addition, we were interested in MA/CA because it is one of the most efficiently cleaved cut sites [18]. We chose to analyze a 12 amino acid region to explore the possibility of selection beyond the 8 positions that have been extensively characterized in complex with PR by crystallography.

To measure fitness landscapes, we engineered a yeast-display approach similar to prior work [4] to quantify cleavage efficiency in high throughput (Figure 1B). We generated plasmids encoding a fusion of Aga2 followed by an HA-tag, a short flexible glycine-rich region, 12 residues of each cut-site (MA/CA, NC/p1, or p1/p6), another short flexible glycine-rich region, and a C-terminal Myc-tag. The Aga2 protein serves to direct the fusion protein to the yeast surface where it is accessible to bulk solution. We used fluorescently labeled antibodies directed at the HA-and myc-tags to distinguish yeast with different levels of cleaved and uncleaved surface-displayed protein. Using yeast displaying only a wildtype (WT) cut-site, we identified conditions that resulted in extensive but not complete cleavage. We reasoned that these conditions would provide high sensitivity by flow cytometry to mutations that either increased or decreased cutting efficiency. We generated site-saturation plasmid libraries at each of the 12 positions for each cut-site and used our flow-cytometry based approach to separate populations based on level of cleavage. We analyzed each FACS population using deep sequencing and used read counts to estimate the frequency of each variant in each sort window. Based on these measurements, we calculated the average fluorescence of each variant. To estimate relative cleavage, we normalized fluorescence estimates to WT, such that 0 represents an uncut site and1 represents the extent of cutting of a WT cut site.

Based on the experimental setup, we expect our readout to be sensitive to changes in both *k*_*cat*_ and *K*_*M*_. Based on the reported range of yeast display efficiency (10^4^-10^6^ molecules per cell) [42] and our conditions that included 10^6^ yeast in a 0.1 mL reaction, we estimate total substrate concentration was 0.16-16 nM. Previously reported *K*_*M*_’s [18, 19] for MA/CA (150 μM), NC/p1 (170 μM) and p1/p6 (1.2 mM) are well above these estimates of substrate concentration and well above the concentration of PR enzyme in our reactions (1 μM). Under these conditions, enzyme theory indicates that both the likelihood of binding PR to substrate (*K*_*S*_) and hydrolysis of bound substrate (*k*_*cat*_) should impact cutting efficiency in our yeast display experiments. Of note, the concentration of the Gag protein containing these cut sites in HIV virions is estimated at about 3 mM [18] . Competition for PR to bind to multiple sites in Gag causes both *k*_*cat*_ and *K*_*S*_ to contribute to the relative rate at which cut-sites in Gag are predicted to be processed in virions [18]. Enzymatic modeling indicates that the changes in cutting efficiency in our yeast display experiments should trend with the efficiency that these cut-sites are processed in virions.

To assess the reproducibility of our approach to measure the protein fitness landscape of a cut-site, we performed a full biological replicate including separate yeast transformations for the MA/CA library (Figure 1C). The relative function of MA/CA cut-site variants was strongly correlated (R^2^=0.97) with the most noticeable distinction between replicates occurring for variants with very low levels of relative cleavage. In addition, independent measurements of WT synonyms at different positions (green symbols in Figure 1C) were tightly clustered. Of note, stop codons in our library generate a displayed protein that lacks the C-terminal myc-tag and thus provide a representation of full cutting (red symbols in Figure 1C). We scaled our measurements of relative function so that 0 represents a fully defective variant, and 1 represents the level of cutting of WT under the experimental conditions. One advantage of bulk competitions such as our yeast display cutting competition is that each variant is exposed to identical conditions including PR concentration, temperature, pH, etc. This internal experimental consistency should contribute to the strong reproducibility that we observed for replicate measurements.

We compared our yeast display estimates of relative cleavage to measurements of cleavage efficiency (*k*_*cat*_*/K*_*M*_) made for a panel of individual cut-site variants for MA/CA and p1/p6 that were analyzed in isolation using a gel readout [40]. As shown in Figure 1D, the relative cleavage rates that we determined correlate roughly with measurements from gel assays for both MA/CA and p1/p6, indicating that both studies capture some shared and fundamental aspects that determine cleavage efficiency. We examined both linear correlations as well as a non-linear correlation with the form of the Michaelis-Menten equation that can account for the a threshold in detection of rapidly cutting variants in the yeast display assay. Of note, the correlation with individual measurements from either fitting approach (R^2^ = 041 or 0.53 for MA/CA) is smaller than the correlation between experimental replicates for MA/CA (R^2^ = 0.97, Fig. 1C), suggesting that distinctions between experimental setups of the yeast display and gel-based assays may impact relative cutting efficiency. Both our yeast display setup as well as the gel-based assays were analyzed using engineered globular protein substrates with triple glycines adjacent to the cut-site to minimize long-range context dependencies. However, the globular proteins differed between each study. Our substrate included Aga2 to facilitate yeast display, while the studies of Potempa et al. [40] utilized GST fused to a truncated MA-CA. The different fused proteins may contribute to variations between the two studies. Of note, amino acid changes that resulted in faster than WT cutting function by yeast display exhibited a wide range of impacts in gel assays (Figure 1D), suggesting that findings of faster than WT cutting should be considered cautiously. The rough overall agreement between our study and those of Potempa et al. [40] indicates that the sequence of the cut-site itself makes a large contribution to cutting efficiency.

### Comparing fitness landscape for the three cut-sites

To search for patterns of mutant impacts on the three cut-sites, we generated heatmap representations (Fig. 2A) of our yeast-display measurements (Supplementary Table S1). To facilitate comparison of variants that cut faster than WT in the heatmaps, an additional scaling of these variants was performed so that the maximum value was identical between each cut-site experiment. From an unfocused perspective the three sites exhibited similar levels of overall selection as all sites had many mutations that severely reduced cutting function (red squares in Fig. 2A) and many mutations that increased cutting function (blue squares in Fig. 2A). The average selection on each amino acid across all positions in all three cut-sites (far right column in Fig. 2A) slightly favored aromatic and negatively charged amino acids, and slightly disfavored amino acids that were either positively charged or polar and uncharged. Aromatic amino acids are enriched at protein-protein interfaces [43, 44] indicating that they have a general propensity to favor binding interactions. The interactions of polar amino acids at protein interfaces tend to be more specific than aromatic interactions because they depend on complementary hydrogen bonding and/or charge [45]. The slight average preference for negatively charged amino acids but not uncharged or positively charged amino acids, suggests the potential for either complementary interactions with positive charge on PR, and/or charge interactions that alter the conformation of our engineered substrate.

**Figure 2.**
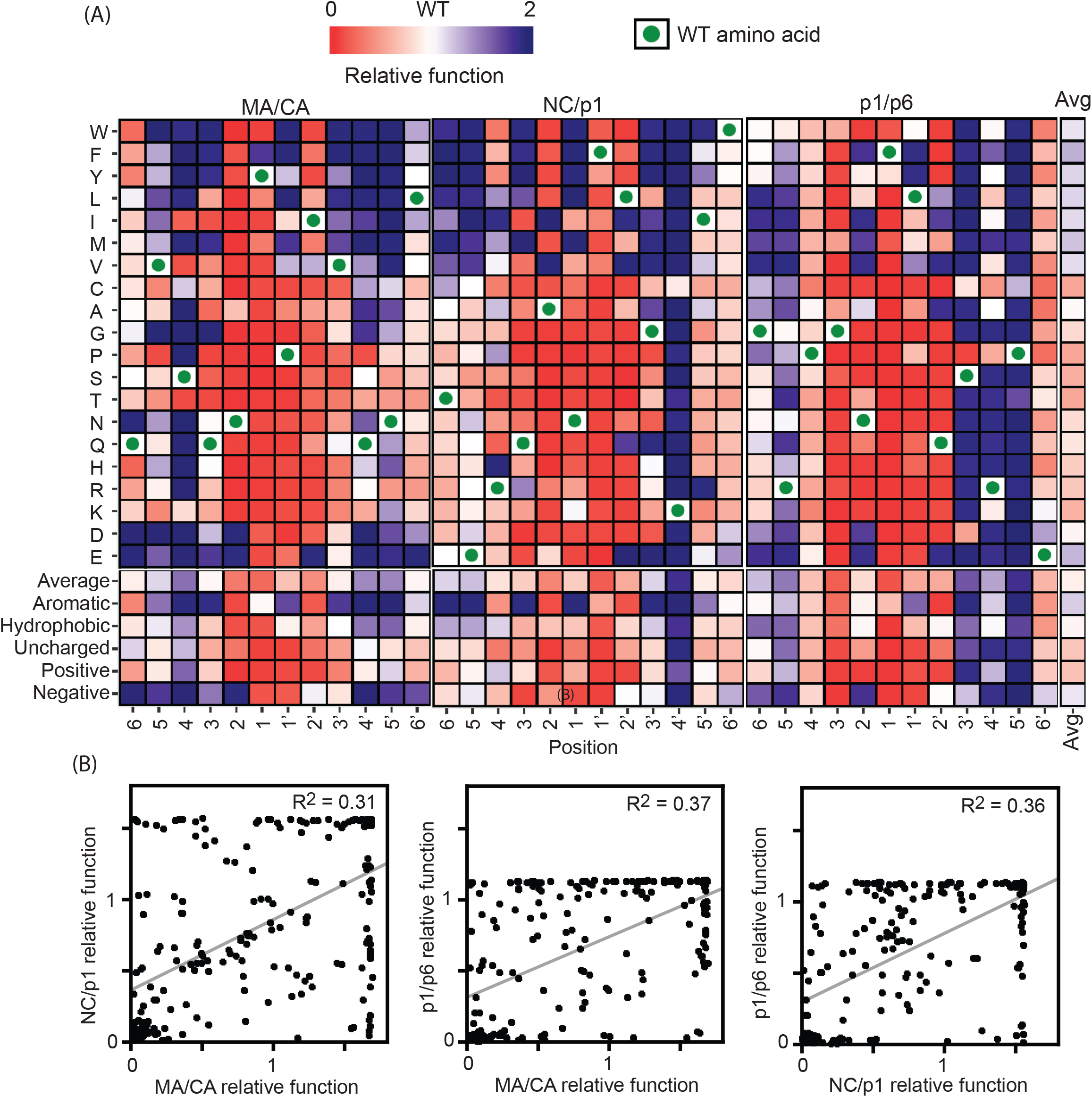
Impacts of amino acid changes in different cut sites. (A) Heatmap representation of yeast-display estimates of cutting function. To facilitate comparison across different substrates, scales were normalized so that the fastest cut variants and slowest cut variants of each cut-site are similar colors. (B) Violin plots comparing rank order of amino acid impacts in different cut-sites across positions 6-6’.

To further investigate the effects of mutations in different cut-sites, we compared the relative function of all mutations in each cut-site (Fig. 2B).We observed correlation coefficients ranging between 0.31 and 0.38, indicating that only a fraction of the observed variation in cutting function was determined by sequence independent impacts of amino acid changes. Amino acid changes that caused very poor cutting in one cut-site tended to have similar impacts in other cut-sites. In contrast, amino acid changes with even modest measured cutting function in one cut-site exhibited a wide variety of effects in other cut-sites. Considering the heatmap (Fig. 2A), we observed stronger and more prevalent decreases in relative function for mutations close to the scissile bond than those further away. Our results also showed that aromatic amino acids could increase relative function in all three cut-sites. However, the location of aromatics that increased activity differed between the cut-sites, indicating some level of sequence-dependence. While visual inspection revealed overall trends, detailed physical and sequence relationships were challenging to extract by visual inspection alone.

### Analysis of common physical underpinnings of fitness effects

To quantitatively search for characteristics that may underly the impacts we observed on relative cutting function, we correlated cutting at each site in each substrate with a set of physical properties of amino acids previously developed for this purpose [46]. The strongest correlation (R^2^=0.75) with cutting effects that we observed was with hydrophobicity at position 6 in the NC/p1 substrate (Fig. 3A). The impacts of mutations at this position in the NC/p1 substrate are modest and span from 0.6 to 1.4. These observations indicate that binding at position 6 in NC/p1 could be mediated by a hydrophobic patch on PR that can accommodate different size and shaped side chains (e.g. a flat and/or flexible hydrophobic surface). Of note, we are not claiming that the correlation in Fig 3A is statistically significant that would require an analyses of multiple hypothesis testing. Rather, we are highlighting how rarely we observed strong correlations between physical properties and relative cutting.

**Figure 3.**
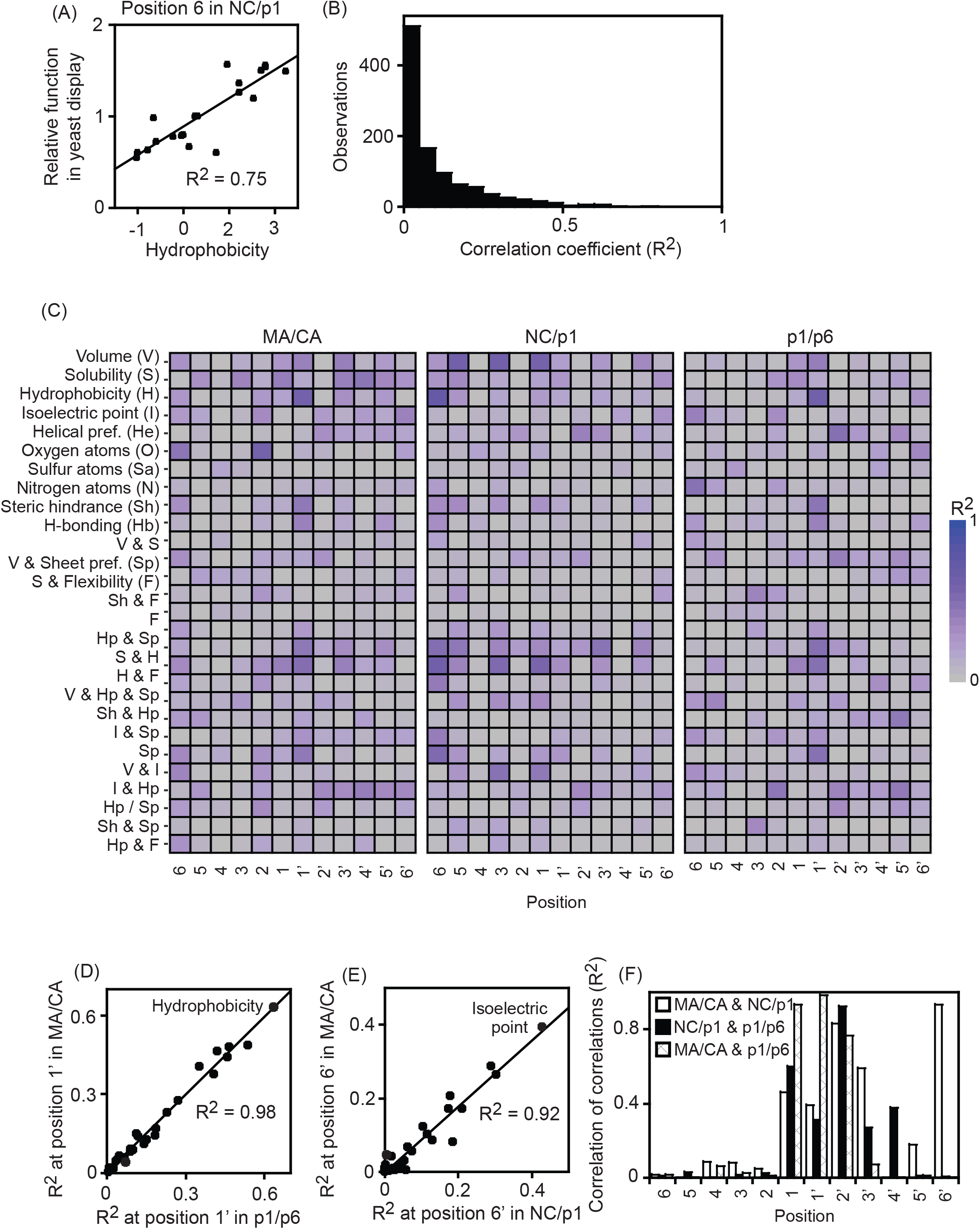
Correlation of physical properties with relative function of positions in different cut-sites. (A) Strongest correlation of property with any position/cut-site was for hydrophobicity at position 6 in NC/p1. (B) Distribution of correlation coefficients between simple physical properties and positions in each cut-site. (C) Heatmap representation of the correlation between physical properties and position/cut-sites. (D&E) Strong correlation of correlations between position 1’ in MA/CA and p1/p6 (D) and position 6’ in MA/CA and NC/p1. (F) Correlation of correlations for the same position in different cut-sites.

Very few physical parameters exhibited even modest correlations (R^2^>0.5) with the observed functional impacts of cut-site mutations at any position/substrate (Fig. 3B). In addition, the strongest correlations between a physical property and a specific site in one substrate generally did not show similar patterns at the same site in the other substrates. For example, while hydrophobicity correlated well with fitness effects at position 6 of the NC/p1 substrate it showed much weaker correlation with cutting effects in MA/CA and p1/p6 substrates. The physical properties that showed the strongest correlation tend to be highly dependent on the sequence background of the cut-site.

Next, we considered the possibility that multiple biophysical parameters with complex interdependencies might underly cutting efficiency. In this case, we expect to see relatively weak correlations between a position and any simple physical property. However, if the complex physical underpinnings at a position were similar across multiple substrates, we would expect to see a correlation of correlations. For example, we would expect similar patterns of physical preferences at a position for multiple substrates. We observed a striking correlation of correlations for a few positions including position 1’ of the MA/CA and p1/p6 substrates (R^2^=0.99, Fig. 3D), and position 1 of MA/CA and p1/p6 (R^2^=0.92, Fig 3E). With the exception of position 6’ in MA/CA and NC/p1, all the strongest correlation of correlations that we observed were close to the scissile bond (Figure 3F). Of note, the high correlations of correlations are skewed towards the C-terminal side of the substrate (p1, p1’, p2’, p3’) for reasons that we do not understand. The correlation of correlations suggests that multiple physical features with complex interdependencies underly cutting at positions close to the scissile bond. Further from the scissile bond, the physical features that mediate cutting efficiency appear to differ between different cut sites.

### Distribution of functional effects (DFE)

To assess the general impacts of mutations at cut-sites, we examined the distribution of functional effects for each cut-site (Figure 4). All three cut-sites showed similar patterns with many amino acid changes causing either large decreases or large increases in cutting relative to WT. For all three cut-sites, there were very few amino acid changes that resulted in WT-like cutting function. Strong defects have been commonly observed in almost all previous analyses of either random or systematic amino acid changes in proteins [47-49]. These general findings are consistent with the cooperative and sensitive nature of the forces that govern protein conformation. Amino acid changes at many/most positions can alter main-chain structure sufficient to severely disrupt function. Most previous analyses of the distribution of effects of amino acid changes in proteins have observed many mutations with little to no effect (WT-like), and very few mutations with increased function relative to WT [47-49], even under stress conditions [50, 51] consistent with natural selection for WT sequences with near-optimal function. In contrast, the distribution of effects that we observe for cut-sites indicates that the WT sequences in our engineered setup are at an intermediate level of potential function. The intermediate level of WT function may have multiple contributing factors that include: balancing natural selection where an intermediate cutting function results in the most effective natural viral fitness; and/or our engineered yeast display setup may artificially alter the way mutations impact cutting function.

**Figure 4.**
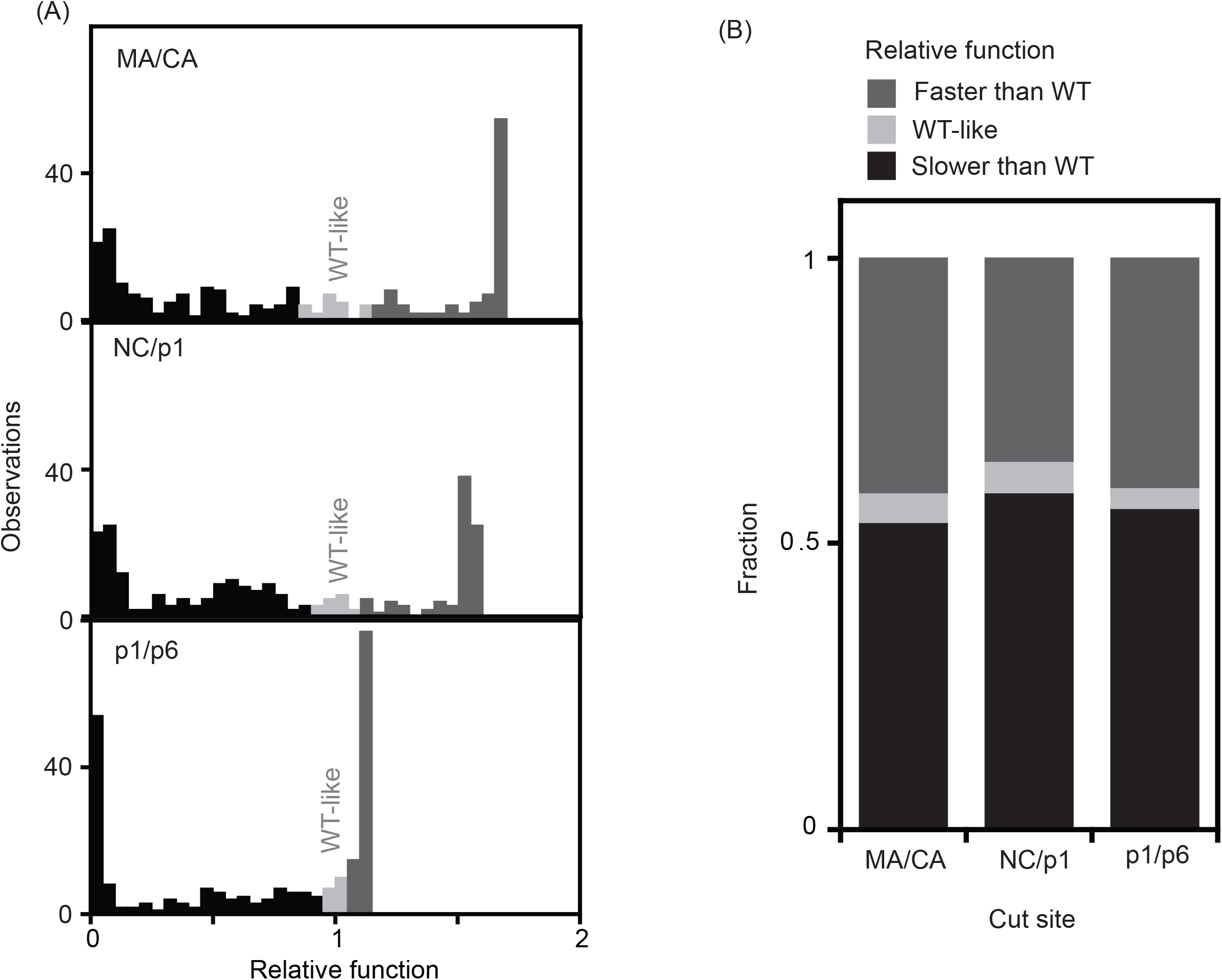
Distribution of mutant effects in cut-sites. (A) Distribution of functional effects measured by yeast-display for point mutants in the MA/CA, NC/p1, and p1/p6 cut sites. Point mutations within two standard deviations of WT synonyms are indicated in light gray, while variants cut slower are in black and variants cut faster are in dark gray. (B) Fraction of WT-like, slower-, and faster-cut variants for each cut-site.

### Variants with increased cutting function

Motivated by the unusually high number of variants that increased cutting function, we analyzed them in further detail (Figure 5). For all three cut sites, amino acid changes that increased function were consistently observed in both experimental replicates (Figure 5A), as expected based on the reproducibility of the yeast display assay. While all three cut sites showed a similar number of variants with increased cutting function, the MA/CA site had the most. We explored the physical properties of MA/CA amino acid changes that increased cutting function. Consistent with the average trends across all three cut sites (Figure 2), the variants with faster cutting of MA/CA contained changes that were predominantly to aromatic and negatively charged amino acids (Figure 5B). The faster cutting amino acid changes in MA/CA tended to occur away from the scissile bond and were most abundant at positions 5, 4, 4’, and 5’ (Figure 5C). The structure of a PR-substrate complex shows that five Lysine and Arginine residues on PR that are in proximity of the last visualized substrate atoms at positions 4 and 4’ (Figure 5D). An electrostatic representation of the surface of PR (Figure 5E) indicates that these lysine and arginine residues generate positive charge that is located adjacent to both the 4 and 4’ positions of substrate. Of note, the structure of the PR dimer is highly symmetrical such that the local environment adjacent to positions 4 and 4’ are nearly identical. Faster cutting of MA/CA substrates in our yeast display assay may be driven by electrostatic complementarity with the positively charged surface region of PR adjacent to positions 4 and 4’ of substrate.

**Figure 5.**
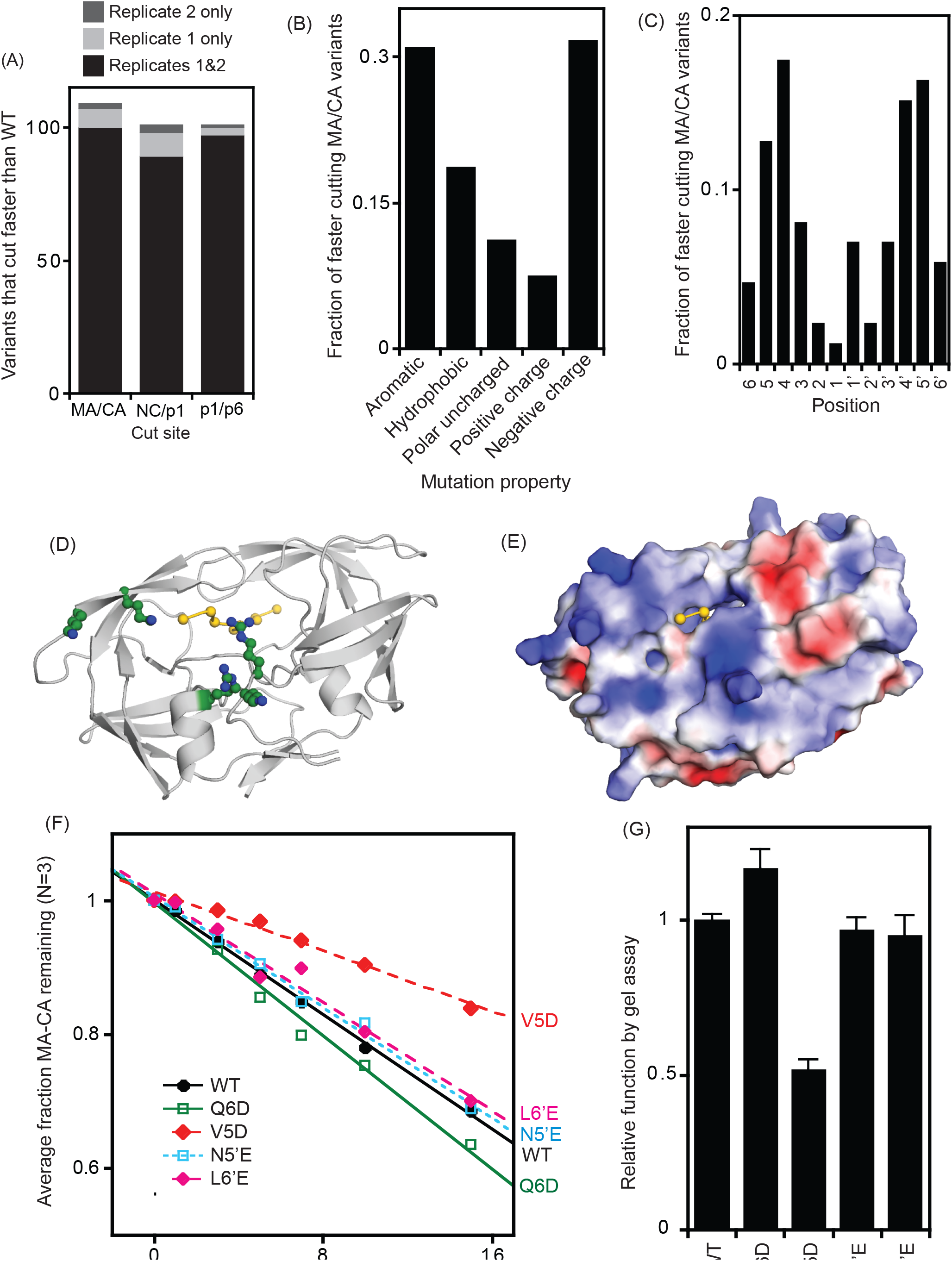
Examining variants that were cut fast by yeast display assay. (A) The number of variants that were cut faster than WT by more than two standard deviations of WT synonyms. (B) The type of amino acids changes in variants of the MA/CA cut site that resulted in faster than WT function. (C) The position of amino acid changes in the MA/CA cut site that were cut faster than WT. (D&E) Structural representation of PR with bound MA/CA peptide substrate based on 1KJ4.PDB [14]. (D) Positions 4-4’ of substrate are shown in yellow with Cα atoms as spheres. Side-chains of lysine and arginine residues of PR that are located in the proximity of the end of the active site are shown in ball and stick representation with carbon atoms colored green. (E) Surface representation indicating electrostatic charge. (F&G) Analysis of the cutting function of a panel of variants in full-length MA-CA substrate using a gel-based assay. Error bars in panel G represent the standard deviation of three assays.

Motivated by the wide range of function seen in gel-based analyses of MA/CA substrates that were fast cutting in our yeast display assay, we investigated an additional panel of MA/CA variants using a gel-based approach. Using purified PR, we measured the cutting of full-length MA-CA protein (Figure 5F&G). The individual variants showed a wide range of cutting in the gel assay using MA-CA as substrate, but none of them were significantly faster than WT. The faster cutting of these variants in the yeast display assay but the WT-like or slower cutting in the gel assay indicate that substrate residues far from the scissile bond can have large impacts on cutting rate. As further discussed in conclusions, our results suggest that the native MA-CA substrate is highly selected to cut rapidly including long-range interactions from sequences far from the scissile bond. By removing these long-range interactions in our engineered yeast display substrates we may have provided artificial opportunities for cut-site mutations to increase cutting rate.

### Comparison of yeast display findings with natural variation in HIV-1

To explore cut-site diversity in circulating HIV-1, we extracted and analyzed sequences of the MA/CA, NC/p1, and p1/p6 cut-sites from the Los Alamos database (http://www.hiv.lanl.gov/). To facilitate the analysis of all three cut sites, we scaled variants that cut faster than WT such that 2 represents full cutting under the experimental conditions. A combined analysis of all three cut sites (Figure 6A) provided similar trends to each site individually (data not shown). Common polymorphisms that were at a frequency of 3×10^−4^ or greater formed two clusters based on the yeast display measures of function, one cluster was faster than WT and one was broad and overlapped with WT (Figure 6A). We chose to consider amino acid changes observed at a frequency of 10^−3^ or greater, corresponding to more than 10 observations, in order to eliminate the majority of null alleles such as stop codons that have been observed in similar HIV-1 sequence datasets [52]. The functional distribution of polymorphisms is similar to the distribution of all mutations with less than a 50% defect in function. In contrast, very few of the polymorphisms showed strong decreases in yeast display function despite this being a common outcome among the comprehensive variants we analyzed. Random sampling of amino acid changes led to a significantly greater number of variants with strong functional defects compared to natural polymorphisms (p<0.01, Figure 6B). These observations indicate that our yeast display assay captures fundamental aspects that determine strong defects in the function of cut-sites and that purifying selection acts to purge these variants from circulating viral populations. In contrast, random sampling led to a similar number of variants with increased function relative to WT according to our yeast-display measurements. These observations are consistent with a lack of selection for faster cutting sites and/or an enrichment of faster cutting measurements caused by artifacts in our yeast display approach.

**Figure 6.**
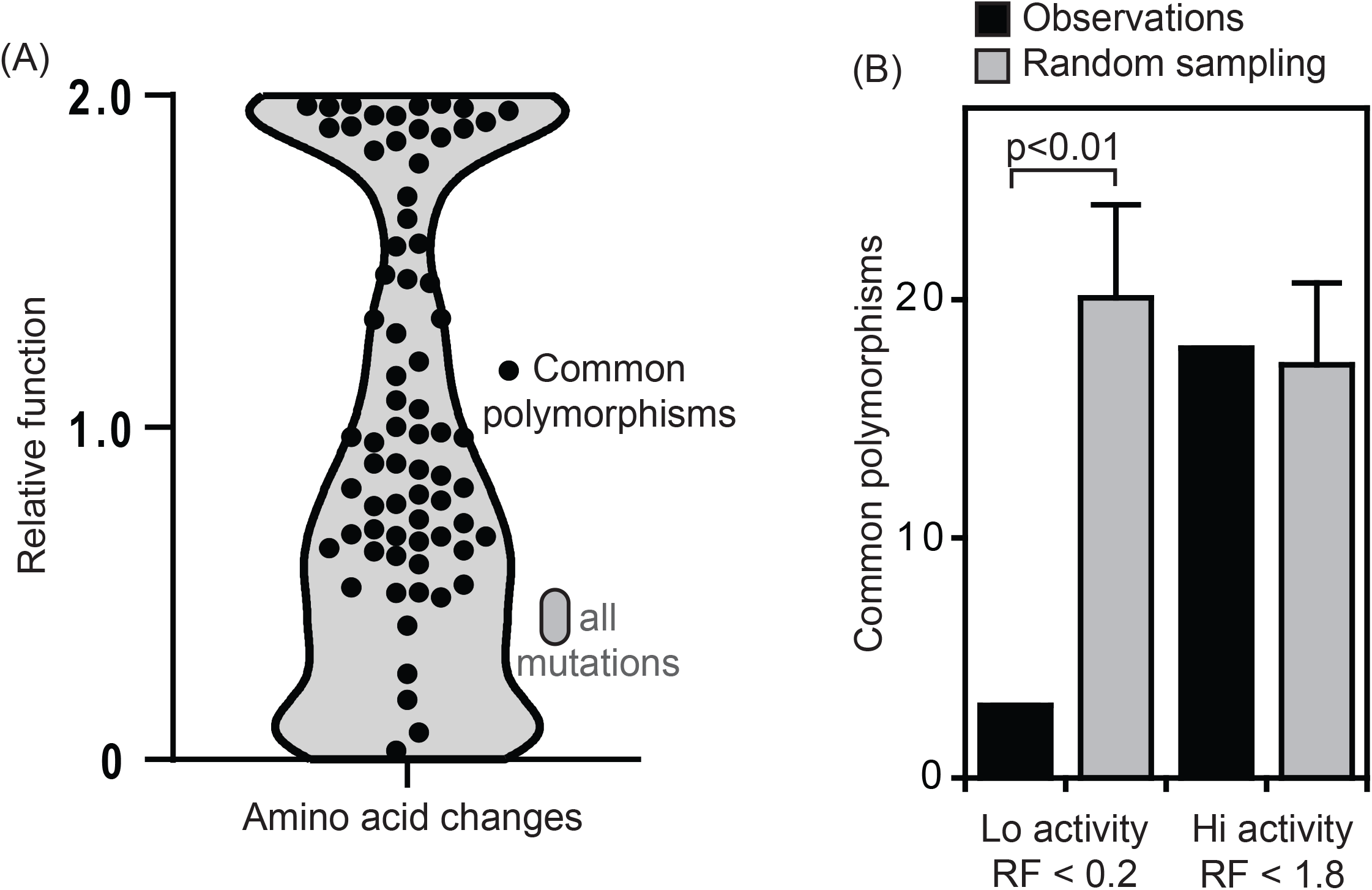
Comparison of functional effects with sequence variation in circulating isolates. (A) Overlay of a violin plot in grey of the functional effects measured by yeast display of all possible point mutations and black dots representing common polymorphisms observed four or more times in a dataset of roughly 12,000 sequenced isolates. (B) Comparison of the number of common polymorphisms with either hi or lo relative function with randomly sampled sets of variants. Error bars indicate the standard deviation from 1,000 random simulations.

## Conclusions

This study highlights how interpreting biochemical analyses in the light of evolution is both important for understanding biology [53] and challenging. One of the biggest challenges in this work was considering potential differences between high-throughput measurements that we made using a yeast-display approach, and the cutting that occurs in viruses and how it relates to fitness. Translating to viral fitness is particularly challenging because cutting in viruses can occur intramolecularly [54], and the relationships between cutting efficiency and fitness remain poorly defined for most cut-sites [18]. Our display approach fused cut sites to a globular protein with cut-sites bracketed by multiple glycines, as similar approaches had been shown to mimic the cutting of biologically relevant substrates [17]. While we observed strong correlation of our yeast-display results with cutting of endogenous substrates for cut-site variants with WT-like or lower function, we found that variants that cut faster than WT by yeast-display had a wide array of function when measured using a full-length substrate (Figures 1&5). In both the yeast display measurements and gel-based analyses, substrate concentrations were well below previously determined Michaelis-Menten values [18], such that both assays should report on impacts on enzyme proficiency (k_cat_/K_m_). In generating conclusions based on our results, we carefully consider that many sites that were cut faster than WT by yeast-display were due to features in the yeast-display substrates that differ from full-length substrates.

Somewhat ironically, this caveat supports one of our main conclusions, that the function of cut-site variants has strong sequence dependencies. Our observations that cut-site variants that were faster than WT by yeast display had WT or lower function when analyzed in a natural substrate context demonstrates that long range sequence dependencies beyond positions 6-6’ contribute to function. Of note, biochemical analyses indicate that regions of substrate far from the scissile bond can contribute to binding to protease [16]. Our findings are consistent with a model where our yeast display construct disrupts or eliminates contacts with protease compared to the natural MA-CA substrate (e.g., Figure 5). In this model (Figure 7), the artificial disruption of contacts in the yeast display setup provides an opportunity for amino acid changes at positions towards the end and past the active site to recoup binding energy with protease. However, in the natural MA/CA substrate, the effect of these amino acid changes has varied impacts because it disrupts the natural interactions of MA/CA with protease.

**Figure 7.**
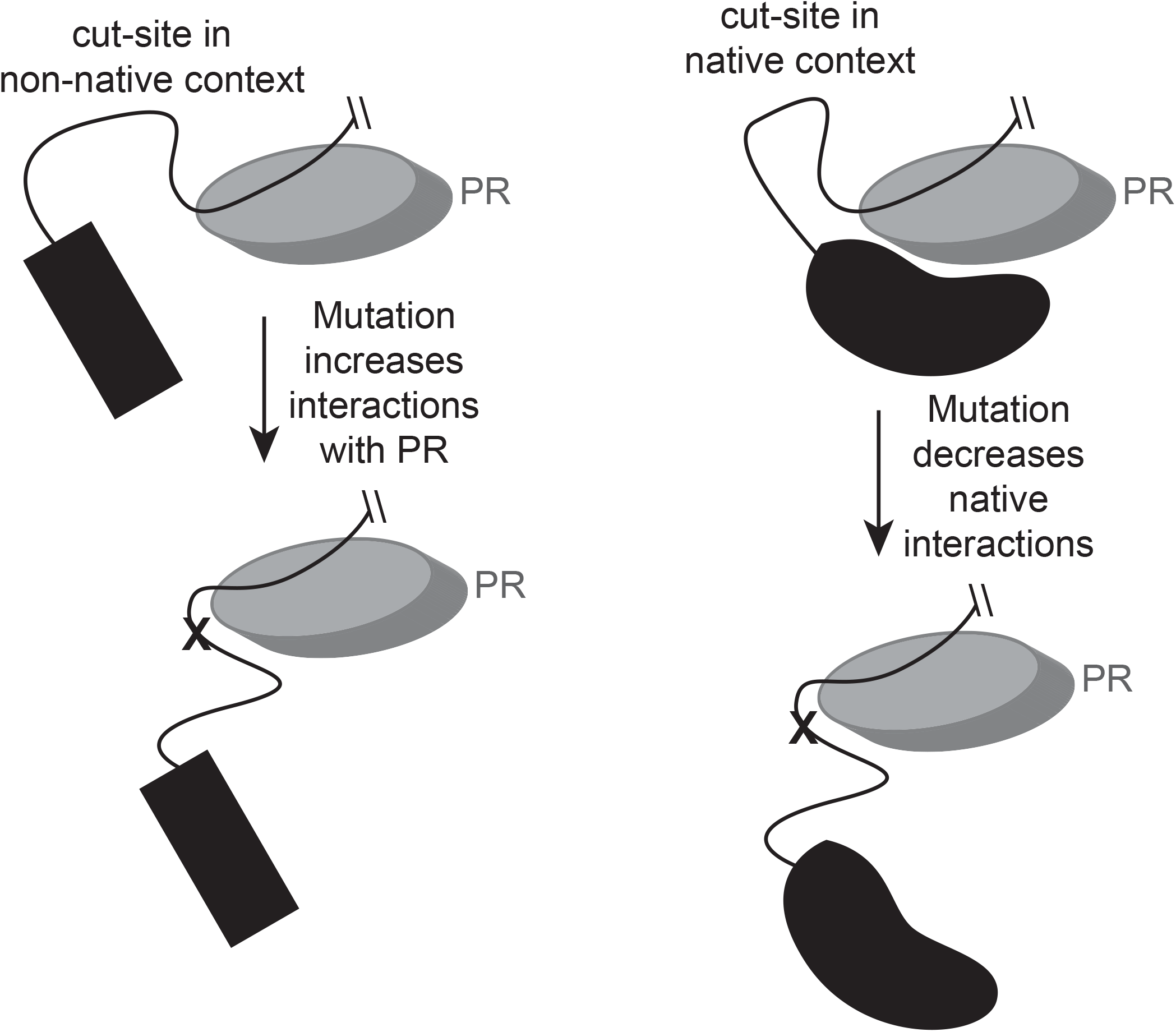
Model of potential distinctions between functional impacts of mutations in engineered cut-sites compared to their native context. Long-range contacts of engineered substrates with PR are unlikely and may provide opportunities for cut-site proximal mutations to increase cutting rate that are not available in native substrates where long-range contacts have been selected.

We also observe strong sequence dependencies throughout most 6-6’ positions in most cut-sites (e.g., Figure 2). Importantly, the comparison between the functional effects of amino acid changes in different cut sites where all measurements were made using yeast display are inherently normalized for the impacts of long-range sequence dependencies because they all are in the same (yet artificial) long-range background. Our analyses indicate that positions 6-3 and 4’-6’ that are distal to the scissile bond had very little to no consistent biophysical underpinnings (Figure 3).

In contrast, we observe consistent but complex physical underpinnings of function at positions adjacent to the scissile bond (Figure 3). We did not find any simple biophysical parameter that correlated strongly with the functional impact of mutations at any position in any cut-site that we tested. However, we do find that the correlations of physical parameters correlates strongest between cut-sites at positions 1,1’, and 2’. This observation suggests that similar physical properties with complex interdependencies mediate cut-site efficiency at positions adjacent to the scissile bond.

This work provides a new and generalizable approach to investigating the relationship between viral proteases and their substrates. We show that long-range and short-range sequence interdependencies are prevalent for HIV-1 cut-site efficiency, but that consistent and complex biophysical properties have large impacts at sites adjacent to the scissile bond. We also demonstrate the importance of careful investigation of experimental setup compared to natural substrates. In future efforts, it will be interesting to examine if similar or distinct physical and sequence-dependent features mediate other viral proteases.

## Methods

### Engineered libraries of cut-site variants

Cut-site variants were engineered in the pCTCON2 yeast-display plasmid [1]. The pCTCON2 plasmid encodes a galactose-inducible promoter driving the expression of a fusion of the Aga2 gene followed by an HA-tag and then a Myc-tag. The fusion protein is directed to the yeast surface by Aga2. We engineered pCTCON2 to encode three glycines and a unique PstI restriction site after the HA-tag and a unique NheI site and three glycines before the Myc-tag. Next, we used cassette ligation to introduce cut-site variants between the PstI and NheI restriction sites. To generate point mutant libraries, we used casettes encoding individual NNK codons (where N indicates a mixture of A,C,G,T and K a mixture of G,T). For each of the MA/CA, NC/p1, and p1/p6 sites, separate cassette ligations were performed with NNK at each of the 12 positions under investigation. Point mutant libraries for each site were generated by mixing libraries from each position (6 to 6’). The resulting plasmids encode Aga2-HA-GGG-CS-GGG-Myc, where CS indicates the cut-site from position 6 to 6’. The wildtype sequence for each cut-site was based on the sequence of NL4-3.

### Yeast surface display

Yeast strain EBY100 [1] was used in all yeast-display experiments. We used the lithium acetate method [55] to introduce plasmids into EBY100 cells. For cut-site libraries, 1 μg of plasmid was used. After transformation with cut-site library plasmids, cells were allowed to recover in 5 mL of synthetic dextrose media at room temperature for 16 hours. Cells were then collected by centrifugation and washed extensively in synthetic dextrose media to remove extracellular plasmid. Cells were then grown in 100 mL of casamino acid (CAA) media with dextrose. CAA-D (CAA with dextrose) media lacks tryptophan, which selects for yeast transformed with the pCTCON2 plasmid that enables tryptophan production. A portion of cells were plated to determine the number of transformed cells. Cells were grown in CAA-D media at 30 ºC until they reached saturation and then mixed with glycerol to a concentration of 25% and frozen at -80 ºC.

Before performing yeast-display assays, a single aliquot for each cleavage site library was thawed and used to inoculate 50 mL of CAA-D media. These cultures were grown in a shaking incubator at 30 °C for 24 hr, and then diluted into 50 mL of fresh CAAD to an OD600 of about 0.1. For the next 6-8h, yeast growth was monitored until the cells had undergone 1-2 doublings. Cells were then collected by centrifugation and washed 3 times with CAA-RG media (CAA media with 1% raffinose and 1% galactose). Cells were resuspended in 50 mL CAA-RG media to an OD600 of about 0.5, and shaken at 30 °C for a further 16 hr. As a control of non-displaying cells, a subset of the culture was grown in CAA-D media.

### Protease treatment, labeling, and FACS of yeast display cells

Yeast display cells were collected by centrifugation. Cells were washed with protease assay buffer (PAB). PAB contained 50 mM sodium acetate at pH 6.0, and 100 mM sodium chloride. 10^6^ cells were resuspended in 100 μL PAB and purified PR was added to 1 μM. Purified HIV-1 PR was a kind gift from the Schiffer lab at UMass Chan Medical School. Samples were incubated in a shaking incubator at 30 ºC for one hour for MA/CA, 30 ºC for 75 minutes for NC/p1, and 30 ºC for 40 minutes for p1/p6 libraries. The protease reaction was stopped by collecting cells by centrifugation and washing them three times in TBSB. A control sample of yeast were left untreated with protease. Following protease treatment, cells were labeled with antibodies to the HA-tag (Alexa488 conjugated, Cell Signaling Technology #2350) and the myc-tag (Alexa647 conjugated, Cell Signaling Technology #2233). Antibody labeling was performed according to the manufacturer’s recommendations using 1:100 dilutions of each antibody and a 1 hour incubation at 23 ºC in TBSB. Labelled cells were washed and suspended in TBSB at a density of 10^6^ cells/mL for FACS.

Cells were sorted on a FACS Aria II cell sorter (BD Biosciences) at the University of Massachusetts Chan Medical School Flow Cytometry Core. Cells untreated with protease were single labeled and used to setup forward-and side-scatter windows that omitted dead and/or aggregated cells. Control cells untreated with protease that were double labeled were used to setup the voltage for robust detection of both labels. Protease treated samples were sorted into four windows: one window encompassing the profile of uncleaved cells, one window encompassing the profile of fully-cleaved cells, and two intermediate windows representing different levels of partial cleavage. A total of at least 150,000 cells were sorted for each window. Sorted cells were grown in 50 mL of CAA-D media on a shaking incubator at 30 °C for 24 hours. The cultures from sorted cells were collected by centrifugation, washed with TBSB and stored as pellets at -80 °C.

### Preparation of DNA and next-generation sequencing

Plasmid DNA was isolated and analyzed essentially as previously described [51]. Briefly, cell pellets were resuspended in buffer P1 (Zymo Research) and treated with zymolyase (Zymo Research) to remove the cell wall. The cells were then lysed with buffer P2 (Zymo Research), neutralized with buffer P3 (Zymo Research), and plasmid was purified using a Zymo-Spin IIN spin column (Zymo Research). Purified samples were then amplified using primers bracketing the cut-site sequence. The forward PCR primer included a P5 sequence and the reverse primer included a P7 sequence in order to facilitate Illumina sequencing. We included a 6-base barcode on the P7-containing primer in order to distinguish samples from different cut-sites and sort windows. PCR products were purified using a Zymo-Spin IIN column following the manufacturers recommendations (Zymo Research). Each PCR product was quantified by qPCR using the Kapa Sybr Fast qPCR Master Mix (Kapa Biosystems) on a Bio-Rad CFX instrument. All PCR products were diluted to 4nM and then pooled for next-generation sequencing. The pooled sample was sequenced on a NextSeq500 sequencer that was kindly made available by the Rando lab at UMass Chan Medical School. A single-end 100 base sequencing run was used to sequence both the cut-site and the indexes that distinguish each sample.

### Estimation of functional effects from sequencing results

We obtained a total of about 3.8×10^8^ sequence reads. Reads were checked for exact match to constant regions between the cut-site and the index. Sequences that passed this quality test and had PHRED scores of 30 or above for all bases encoding the cut-site were then parsed into separate files based on the index read. The sequence of the cut-site in these reads were then analyzed and the counts of all individual amino acid changes tabulated. In each sort window, we calculated the frequency of each variant (*F*_*i*_) as the sequence counts of that variant divided by the total sequencing counts for the window. In order to account for the asymmetric distribution of variants across the FACS windows, we calculated the relative window size (*W*_*j*_) as the fraction of cells from the protease treated library sample in each window. We then quantified the activity (*A*) of each variant using an approach similar to a center of mass calculation using the following equation:

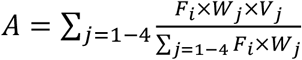

Where *V*_*j*_ is the value associated with each window. We used values of 1-4 for *Vj* representing the window with the least cutting to the window with the most cutting. These activity estimates range from 1 for a variant that was uncut to 4 for a variant that was fully cut. To facilitate comparison between the effects of amino acid changes in each cut-site, we estimated relative function (RF) by normalizing the activity measurements with linear transformations such that 0 represents an uncut variant (based on the average of the five most defective variants in each library) and 1 represents the wildtype cut-site. For the heatmap analyses in Figure 2 and the comparison to diversity in circulating variants in Figure 6, we performed a second linear normalization on variants with faster than wildtype RF such that the average stop codon had a value of 2. This normalization was done to account for varied sensitivity to detect faster cutting variants due to distinctions in the fraction of cutting of the wildtype for each cut-site library.

### Gel-based analyses of the cutting of full-length MA/CA variants

Full-length MA-CA was cloned into pET21 with an in-frame 6xHis tag at the C-terminus. Cut-site variants were generated by site-directed mutagenesis. Plasmid variants were transformed into BLR(DE3) pRIL bacteria that were grown in 2xYT media to an OD600=0.8 and induced with 1 mM IPTG for 3 hours. Cells were collected by centrifugation and stored at -80 ºC. Cell pellets were thawed and resuspended in 25 mM HEPES pH 7.5 with 500 mM sodium chloride. Cells were lysed by sonication and cell debris pelleted by centrifugation. Soluble MA-CA protein was purified from the supernatant by nickel affinity chromatography. Purified protein was dialyzed into 25 mM HEPES pH 7.5 with 10 mM sodium chloride and further purified by anion exchange chromatography. Purified protein was dialyzed into 25 mM HEPES pH 7.5 with 10 mM sodium chloride. The concentration of MA-CA protein was determined based on absorbance at 280 nm. Cutting assays were performed at 25 ºC in 50 mM sodium acetate at pH 6 with 100 mM sodium chloride. Cutting assays contained 250 nM of purified PR and 10 μM of MA-CA variants.

Timepoints samples were taken and the reaction stopped by the addition of 2% SDS and heating to 95 ºC for 2 minutes. Timepoint samples were run on a denaturing polyacrylamide gel and stained with Coomassie brilliant blue. The density of the MA-CA band was quantified using an Amersham Imager 600. All gel-based cutting assays were performed in triplicate.

### Analysis of cut-site variation in sequenced isolates

We analyzed the cleavage site sequences of MA/CA, NC/p1 and p1/p6 in roughly 12,000 HIV-1 subtype-B isolates in the Los Alamos HIV database (http://hiv-web.lanl.gov). The majority of sequenced isolates (roughly 11,000) were from drug-treated individuals. Variants were defined as changes in amino acid with respect to the sequence of NL4-3. We identified 71 amino acid variations across all three cut sites that we analyzed (MA/CA, NC/p1, p1/p6) that were observed at a frequency of 10^−3^ or greater. Roughly, equivalent percentage of these 71 amino acid changes were from each cut-site (38% for MA/CA, 29% for NC/p1, and 33% for p1/p6). Bootstrap analyses were performed by randomly sampling 71 amino acid changes from the combined set of functional scores for all three cut-sites with faster cutting sites rescaled so that 2 represents full cutting under the experimental conditions (Figure 6A). One thousand random simulations were performed and the average and standard deviations of sites with high functional scores (>1.8) and low functional scores (<0.2) were shown in Figure 6B.

## Acknowledgements

This study benefited tremendously from useful input and discussions with Prof. Celia Schiffer and Dr. Julia Flynn. This work was supported by funds from grant R01GM112844 from the National Institutes of Health to D.N.B. The authors declare that they have no conflicts of interest with this work.

